# Low-troposphere microbial communities differ between dry-air and rainfall but do not show strong seasonal patterns in Metro Atlanta, Southeast USA

**DOI:** 10.1101/2023.06.09.544356

**Authors:** Lizbeth Davila-Santiago, Casey Erb, Laura Hyesung Yang, Johanna Hall, Arnaldo Negron, Isabelle D’amico, Janet K. Hatt, Konstantinos T. Konstantinidis

## Abstract

The composition and seasonal patterns of airborne bacterial and fungal communities and how these are affected by atmospheric conditions (e.g., dry vs. rain), origin of air masses, and presence of air pollutants remain understudied, despite their obvious importance for public health. To provide insights into these questions, monthly dry air and rain samples were collected at the Environmental Science and Technology building rooftop on Georgia Tech’s campus (Metro Atlanta) between June 2017 and November 2019. The sampling included the remnants of Hurricane Irma and a Saharan dust event in 2020. Amplicon sequencing of the V4 region of the 16S rRNA gene and the fungal nuclear ribosomal internal transcribed spacer (ITS) region showed that spore-forming bacteria and widespread fungi were enriched in dry samples, while photosynthetic bacteria and wood-decaying fungi were more abundant in rain samples, demonstrating the effect of sample type on bioaerosol composition. Further, higher relative abundance of fungal human pathogens and allergens were identified in the dry-air and Saharan dust samples, including *Alternaria alternata* and *Cladosporium cladosporioides*. Bacterial and fungal species richness and composition appeared to be relatively consistent between seasons for both sample types. Accordingly, sample type and seasonality explained ∼14% and ∼8.5% of the microbial diversity between samples, respectively, while presence of air pollutants and three-day back trajectory data were not significant. Collectively, our data indicates that dry air might represent a higher public health risk and provides a reference point for the long-term monitoring of airborne microbial communities in an urban Southeast US setting.

**Importance:** In the atmosphere, or in the air we breathe, bioaerosols are always present. Bioaerosols are biological particles (alive or dead) suspended in the air, being bacteria and fungi the most abundant. In addition, bioaerosols can potentially contribute to weather/climate patterns. Although and unfortunately, a clear biodiversity pattern from different atmospheric events (air, rain, snow, etc.) remains to be discovered, especially in urban areas, where bioaerosols can also have implications for public health. The role of airborne microbes and their diversity patterns in the atmosphere constitutes a significant gap in our understanding of their interactions with health, climate, and other ecosystems compared to other environments. Our research provides the first reference point for long-term monitoring of airborne microbial communities in an urban Southeast US setting. This research contributes novel knowledge about public health and insights for integrating biological information into weather and climate prediction models.

## Introduction

Microorganisms are known to inhabit most of the ecosystems on Earth, including the atmosphere. However, the atmosphere, although known to support microbial life, has received little attention for its biology due to the traditional focus on its physical and chemical properties (Polymenakou, 2012). Bioaerosols are biological particles (whole cells or cellular components) such as bacteria, fungi, pollen, and viruses (Lacey & Dutkiewicz, 1994; Fröhlich-Nowoisky et al., 2016) suspended in the atmosphere. Bacterial and fungal cells are the most abundant bioaerosols with concentrations near the ground ranging from 10^2^ to 10^6^ cells/m^3^ (Bauer et al., 2002; Bowers et al., 2012; DeLeon-Rodriguez et al., 2013) and 10^3^ to 10^6^ cells/m^3^ (Bauer et al., 2002; Fierer et al., 2008), respectively. Bioaerosols originate from a large diversity of sources, including soils, plants, animals, waterbodies, and human activity (Fröhlich-Nowoisky et al., 2016; Ruiz-Gil et al., 2020; Xie et al., 2020). In addition, bioaerosols have been reported to remain airborne for long periods of time (even several weeks) as well as to travel long distances (Mayol et al., 2017; Tignat-Perrier et al., 2019; Smith et al., 2013; Uetake et al., 2019; Uetake et al., 2020). Therefore, bioaerosols can potentially impact air quality and public health, including the health of animals and plants, over long distances. Finally, bioaerosols could play a role in cloud formation and precipitation by serving as cloud condensation nuclei (CCN) or ice nuclei (IN), directly affecting the physics and chemistry of the atmosphere and the hydrological cycle (Huffman et al., 2013; Morris et al., 2014; Amato et al., 2015; Fröhlich-Nowoisky et al., 2016b).

Bioaerosols are eventually removed from the atmosphere through dry (sedimentation) or wet (rain or snow) deposition. Dry and wet depositions are both important mechanisms for controlling air pollution and removing particulate matter (PM) and bioaerosols (Shannigrahi et al., 2005; Wu et al., 2018). Wet deposition is reportedly more important in removing bioaerosols from the atmosphere (Fröhlich-Nowoisky et al., 2016; Joung et al., 2017), but dry deposition has been proposed to be more relevant for local air quality and health effects (Fröhlich-Nowoisky et al., 2016).

Studies that evaluated the influence of rainfall on microbial community composition primarily took aerosol samples during non-rainy vs rainy days, as opposed to directly sampling rain. These studies were also only conducted in forested and marine environments (Huffman et al., 2013; Evans et al., 2019). Studies that evaluated microbial composition between dry-air (on non-rainy days) and rain samples were either only focused on fungal communities (Woo et al., 2018), or were conducted at high elevation mountains and remote places (Els et al., 2019; Triadó-Margarit et al., 2019). These studies revealed higher fungal species richness in wet deposition (Woo et al., 2018). However, the relevance of these findings for metropolitan and/or sea-level areas remain unknown. In addition, most of the previously mentioned studies that investigated dry and wet deposition did not evaluate seasonality patterns, while other studies have shown a seasonal pattern in airborne microbial communities in (mostly) dry samples. For instance, one study analyzed rain and snow samples from high elevation of a mountain in the Central Pyrenees, Spain and found substantial differences in airborne microbes between summer and winter seasons (Cáliz et al., 2018). Therefore, studies of dry-air and rain samples collected at the same location and across seasons are lacking, especially in urban settings in the Southeast US, but are critical for obtaining a more complete view of airborne microbial diversity patterns and their relevance for public health. In the present study, we evaluated the composition and diversity of bacterial and fungal communities in dry-air and rain samples, accounting for seasonality and other environmental factors in Atlanta, Georgia, US. We report on samples taken every month for two consecutive years, providing a unique time-series both in the frequency of samples as well as the geographic location. Notably, the broader Atlanta area is characterized by both urban influence and frequent continental and oceanic air transport and is surrounded by deciduous forests. Although the city of Atlanta has been studied for bioaerosols previously, research has been solely focused on quantification of the total concentration of bioaerosols during the spring season or detection of pathogens near wastewater treatment plants (Negron et al., 2020; Ginn et al., 2021). We hypothesized that seasonality plays a role in shaping bacterial and fungal composition, in addition to the effect of other environmental factors, such as pollution and air transport.

## Methods

### Rain sample collection

Rainwater samples were collected on the rooftop of the Ford Environmental Science & Technology Building (approximately 12 m above ground level) on the Georgia Institute of Technology (Georgia Tech) campus in Atlanta, Georgia between June 2017 through November 2019. Samples were collected approximately 3 – 4 times per month on random days or days selected based on weather patterns. For every rain event sampled, the rainwater was funneled into three sterilized 1L glass bottles and later pooled together during the filtration step. Funnels were cleaned and sterilized by: (i) scrubbing with 1% Alconox, (ii) rinsing with (1 L volumes and 3 times) with DI water, (iii) spraying with 20% bleach, (iv) rinsing again with sterile DI water. Once on the rooftop, each funnel was cleaned with 70% alcohol just before use in sampling. The duration of each rainwater sampling was variable from sample to sample depending on rain duration and intensity. Rainwater collection lasted until the rain event stopped or the sample bottles overflowing. After sampling, the collected rainwater was taken to the laboratory and immediately filtered through a UV-treated 0.22 µm pore size filter (nitrocellulose 47mm). For each rainwater sample, a second filter was used as a handling blank. Filters were stored at −80°C until used for DNA extraction.

### Dry-air sample collection

Dry-air samples were collected the same months at the same location as the rain samples. Dry-air sampling days were selected when the weather forecast did not predict rain events for at least 24 hrs. Dry samples were also taken during a Saharan dust cloud event in the summer of 2020. Samples were collected using a Thermo Scientific MFC-PM10 High Volume Air Sampler for a duration of 24 h per sample. The instrument uses a mass flow-controlled system for sampling airborne particulate matter less than 10 μm in diameter. Flowrate for the sampler is approximately 1,132 L/min. Dry samples were collected on an 8 x 10 in Quartz filter, which was previously baked at 500°C for 48 hrs. After sampling, the filter was transported to the laboratory and one quarter (¼) of the filter was cut into strips with sterile scissors and forceps, placed inside a 50 mL centrifuge tube, and stored at −80°C. This filter sample was later used for DNA extraction, while the remaining three quarters (¾) of the filter is stored at −80°C in sterile Whirl-Pak bags (123 oz) as an archived sample.

Before DNA extraction, the filter sample is washed in 1X phosphate-buffered saline (PBS) (pH = 7.6) for detachment of microbial cells. Forty milliliters of 1X PBS buffer filter sterilized through a 0.22 µm filter was added to a 50 mL centrifuge tube containing the filter sample and vortexed for 3 min at 1600 rpm. The filter strips were the transferred aseptically to a new 50mL centrifuge tube and the wash repeated. The 80 mL of collected cells was then filtered through a UV-treated 0.22 µm pore size filter (nitrocellulose 47mm) and used for DNA extraction.

### Metadata records

In addition to type of sample, season, month, year, time of day (AM or PM), volume of air/rain collected, and weather conditions before sampling (type of day; dry or rain), a variety of atmospheric conditions were recorded. Temperature, relative humidity, rain gauge, rain rate, solar radiation, wind gust, and other factors for each sampling day were retrieved from the WeatherSTEM station at Georgia Tech (https://gatech.weatherstem.com/). Data about pollen and mold for each sampling day were retrieved from the Atlanta Allergy and Asthma website (https://www.atlantaallergy.com). Air quality data such as PM2.5 and PM10 were retrieved from the AirNow website (https://www.airnow.gov), while other air pollutants regulated by the Clean Air Act by the US EPA, which include nitrogen dioxide (NO_2_), carbon monoxide (CO), sulfur dioxide (SO_2_), and Ozone (O_3_), were retrieved from the US EPA website (https://www.epa.gov/outdoor-air-quality-data). Finally, back trajectory analysis was conducted with the Hybrid Single-Particle Lagrangian Integrated Trajectory model (HYSPLIT) by NOAA’s Air Resources Laboratory to determine the cardinal and intercardinal directions where the air mass originated 72 h before the sampling date. The HYSPLIT model is commonly used for atmospheric trajectory and dispersion calculations (Stein et al., 2015). Representative samples for back trajectory analysis with HYSPLIT models are shown in Fig. S1.

### DNA extraction

DNA from biomass collected on filters were extracted with a phenol-chloroform protocol in a sterile 2 mL O-Ring screw cap microtube. First, approximately 0.20 g of each 0.1mm glass and 0.5 mm ziconia/silica beads were added to the sterile microtube. Next, RNase to 100 mg/mL and lysozyme to 1.15 mg/mL final concentration were added and incubated at 37°C for 30 min with mixing. After incubation, 40 uL of Proteinase K solution (10 mg/ml) and 60 uL 10% SDS were added to the microtube and incubated again at 55°C for 2 hours, with mixing. Subsequently, 600 uL of phenol:chloroform:isoamyl alcohol (25:24:1) was added, the tube vortexed vigorously for a few seconds and incubated at 65°C for 5 min. Then, the tube was centrifuged at 10,000 rpm for 5 min and the upper aqueous phase was transferred to a UV-treated 1.7 mL Lo-bind microtube. An equal volume of chloroform was added, and the mixture was inverted to mix for 10 s. A second spin at 10,000 rpm for 5 min was performed and the aqueous phase was transferred to a fresh UV-treated microtube. The DNA was precipitated by addition of 1/10 volume of sodium acetate (3M), 1 uL of glycogen, 2.5x volumes of ice cold 100% molecular-grade ethanol thoroughly mixed and incubated at −20°C overnight. After precipitation, the DNA was centrifuged at 13,000 rpm for 30 min at 4°C to obtain a pellet. The pellet was washed 2 times with 1 mL of cold 70% ethanol. Finally, the pellet was dried for 12 min with a SpeedVac concentrator (no heat) and 35 uL of sterile EB buffer (10 mM-Tris-Cl; pH = 8.5) was added and mixed well to dissolve the pellet. DNA was evaluated with Nanodrop for purity values and assessed by Qubit 1X HS dsDNA assay for accurate DNA quantification.

### PCR amplification of the V4 region of the 16S rRNA gene and fungal ITS2

PCR reactions for 16S rRNA gene V4 amplicon sequencing, based on the protocol described previously (Kozich et al., 2013), were performed on the extracted DNA. First, PCR tubes (0.2 mL) and micropipets were sterilized under UV-light for 20 min. Unique primer combinations were prepared for each sample in nuclease-free water for a total concentration of 5 uM per primer. For master mix preparation, the following solutions were mixed in a 1.7 mL tube for each 25 uL PCR reaction intended: 17.25 uL of nuclease-free water, 2.5 uL of AccuPrime 10x Reaction Mix, 1.25 uL of BSA (10 ng/uL) and 0.25 uL of AccuPrime Pfx DNA Polymerase. The PCR reaction tubes were prepared with reagents added in the following order: 21.25 uL of master mix, 2.5 uL of a unique primer combination, and 1.25 uL of DNA template (1:10 dilution of DNA in EB buffer) and mixed well by pipetting. Reactions were amplified as follows: initial denaturation at 95°C for 2 min, then 30 cycles of denaturation at 94°C for 30 s, annealing at 55°C for 30 s, elongation at 72°C for 1 min, and a final extension step at 72°C for 6 min. For fungal ITS amplification degenerate primers targeting the ITS region (Bengtsson-Palme et al., 2013) were modified using the method as described for amplification of the V4 region of 16S rRNA (Kozich et al., 2013). PCR settings used were an initial denaturation at 92°C for 2 min followed by 25 cycles of denaturation at 95°C for 30 s, annealing at 52°C for 30 s, elongation at 72°C for 1 min, and a final extension step at 72°C for 6 min. A negative control with sterile, nuclease-free water as template was included in all PCR runs. Agarose gel electrophoresis was run after every PCR and each sample analyzed for successful amplification (expected amplicon size for bacterial V4 16S rRNA ∼ 385 bp; fungal ITS ∼ 550 bp). Duplicate samples were pooled and cleaned using 35 uL of SPRI beads from Beckman Coulter® Life Sciences.

### Sequence analysis of amplicons

After amplification, PCR products were sequenced on an Illumina MiSeq platform for 500 cycles (250 bp paired end run) at the Molecular Evolution Core, Georgia Institute of Technology. Raw sequences were uploaded to QIIME2 2021.4 version (Bolyen et al., 2019) for sequence analysis for a total of 72 field and 6 control samples. Demultiplexed samples were denoised using the DADA2 (Divisive Amplicon Denoising Algorithm) (Callahan et al., 2016), with command “Qiime dada2 denoise-paired” integrated in QIIME2 2021.4 version tool. Based on DADA2 quality control results, reverse reads were truncated to 200 bp for downstream analysis. The final product is a table (and corresponding sequences) of the exact amplicon sequence variants (ASV) present in the samples, which is a higher-resolution analogue of the traditional OTU table and includes how many times each ASV was observed in each sample (relative abundance). We also performed OTU clustering at a 97% cutoff within QIIME2 to compare ASVs and OTUs data. The taxonomic profile was similar for both methods, but community diversity was slightly higher in ASVs than OTUs, as expected. We continued with the ASVs approach since it can provide an advantage for a more accurate identification of microbes due to its higher resolution.

Using QIIME2, the recovered ASVs were searched against the Silva SSU database 138 (Quast et al., 2013) for taxonomy identification of bacteria and against the UNITE database version 8.3 (Abarenkov et al., 2021) for taxonomy identification of fungi. Sequences classified as *Escherichia-Shigella*, chloroplast, and mitochondria were removed from all samples. *Escherichia-Shigella* was removed for its significant abundance in control samples. Samples having 5000 sequences or less after removing the *Escherichia-Shigella*, chloroplast, and mitochondria sequences were discarded from further analysis. In addition, all the ASVs identified in the six control samples of the bacterial dataset and the single control sample from the fungal dataset were removed from field samples (if present) as potential contaminants, similar to what was performed previously by Evans and colleagues (2019) and Kobziar and colleagues (2022). Alpha-diversity (chao1 richness index and Pielou’s evenness index) and beta-diversity (Bray-Curtis dissimilarity) analyses were performed by QIIME2 and plotted using RStudio (v. 2021.9.0.351) with the ggplot2 and vegan packages, respectively.

To further validate results using the QIIME2 pipeline and quantitatively assess how our metadata explains the variance among the samples, permutational multivariate analysis of variance using distance matrices was performed with the Adonis function within the vegan package in RStudio. Adonis is a function for the analysis and partitioning of sums of squares using dissimilarities and is based on the algorithm of Anderson (2001). The Bray-Curtis matrix exported from QIIME2 was used for the Adonis analysis, which evaluates how similar communities are when considering specific conditions or treatments and accounts for relative abundance of ASVs. The Bray-Curtis distance matrix was also used to evaluate how similar communities are when considering the effects of sample type, season, and other measured environmental factors. Finally, non-metric multidimensional scaling (NMDS) plots using the Bray-Curtis distance matrix were generated in RStudio for visualization of results with the vegan package.

Linear discriminant analysis (LDA) effect size (LEfSe) analysis was performed to identify significant differences in the abundances of bacterial and fungal ASVs for each sample type, with a threshold of 3.0. LEfSe will determine which ASVs are most likely to explain differences between our sample groups (Segata et al., 2011).

## Results

### Sample description and metadata

Outdoor bioaerosol samples are typically low in biomass, which presents a series of challenges during sample processing and DNA extraction steps (Salter et al., 2014; Weiss et al., 2014; Eisenhofer et al., 2019). The DNA concentration of the samples obtained in this study ranged from 0.067 ng/μL to 21.5 ng/μL, with samples from the spring season showing the highest DNA concentrations (Table S1). All available metadata for each sample are reported in Table S1.

DNA was extracted from 91 bioaerosol field samples (42 dry air; 43 rainwater; 6 Saharan Dust) collected from June 2017 through November 2019 including rain samples from the remnants of Hurricane Irma and samples from the 2020 Godzilla Saharan dust storm event (Francis et al., 2022). In addition, 50 control samples were subjected to DNA extraction to address any potential contamination resulting from filter handling, DNA extraction, PBS buffer, or aerial cross contamination. Successful amplicon sequencing was obtained for 73 field and 6 control samples for the 16S rRNA gene, while 73 field and 1 control sample yielded amplicons for the fungal ITS marker. A total of 70 samples had both 16S rRNA gene and ITS amplicon datasets. Seasonal breakdown of samples for the bacterial dataset were 18 winter, 16 spring, 21 summer, and 18 fall, while for the ITS dataset, sequenced samples were composed of 15 winter, 17 spring, 23 summer, and 18 fall samples.

### Bacterial/V4-region sequence statistics and assessment of contamination

The average number of raw sequences per sample for the bacterial dataset was 57,897 for field samples, while the number of raw sequences for control samples was 9,866. After denoising with DADA2, 62% of the sequences, on average, were retained (Fig. S2). Following ASV identification and classification, chloroplast and mitochondria sequences were identified in all the field samples, most abundantly in late winter and spring seasons (blooming season in Atlanta). These sequences were absent from the control samples, except for sample RC021919 (Fig. S3), suggesting that our sampling acquisition and processing pipeline was robust against contamination. Only 12% of control samples produced amplicons while 86% of field samples were amplified and successfully sequenced, supporting the proficiency of our protocol. The presence of chloroplast and mitochondria in bioaerosol samples is not uncommon (Fahlgren et al., 2011; Evans et al., 2019; Petroselli et al., 2021; Tang et al., 2022), therefore our bacterial dataset was expected to contain some sequences derived from the genomes of these organelles. These results also agree with higher plant activity and blooms in February-March for the Southeast US compared to other months. To be conservative in our conclusions, ASVs (n = 196) found in any of the six sequenced controls, such as *Escherichia-Shigella,* were removed from the field samples prior to any subsequent analysis. On average, only 1.8% of sequences per field sample were removed by this step, which suggests a low level of contamination of the field samples. Distribution of all ASVs detected in control samples across the field samples as well as sequencing depth for each sample are shown in Fig S4. In addition, taxonomic analysis of control and field samples suggested a very different bacterial composition between these two groups (Fig. S5). Thus, potential contamination in control and field samples might have come from the molecular reagents or aerosol contamination in the laboratory during sample processing but was overall minimal.

After denoising sequences, removing ASVs found in control samples, and removing ASVs belonging to chloroplast and mitochondria, 11 field samples (6 dry, 5-rain), mostly from spring and high-pollen seasons, were removed from further analysis since the number of remaining sequences was below 5,000 for these samples vs. an average of 20,000 in the remaining field samples. The 11 field samples removed from the dataset had an average of only 7% recovery after the denoising step, indicating that they were compromised and/or extremely low biomass microbial samples, likely due to high concentrations of pollen as mentioned above. Finally, for the remaining field samples (n = 61), the average percentage of recovery was 39% (Fig. S2) from the original input sequences with a final average of 22,256 sequences per sample. Most of the sequencing reads removed were plant and mitochondrial sequences, especially in the spring season samples.

### Fungal/ITS sequence statistics and ASV filtering

The average number of raw fungal sequences per sample was 22,838 for field samples. After denoising, there was an average recovery of 59% (Fig. S6). Removal of chloroplast and mitochondria was not necessary in the fungal dataset since the ITS primers do not amplify chloroplast and mitochondrial DNA. Thirty-six ASVs were removed from the field samples as they were detected in the single control that was successfully sequenced out of two control samples taken in total while processing samples for ITS analysis. More control samples were taken during sampling, but any DNA present in these samples was consumed during V4-region amplification and thus, only two samples were available for ITS amplification. The control sample (RC0415) had only 1783 sequences (Fig. S7). After denoising and removing control ASVs, the average number of sequences for the field samples was 11,829 with an average recovery of 51% from the original sequences input (Fig. S6). In addition, taxonomic analysis revealed a very different fungal composition between control and field samples (Fig. S8). After quality control, the fungal dataset had a total of 72 samples, with 60 samples overlapping between the bacterial and fungal datasets (Table S1).

### Microbial richness is similar between dry-air and rain in Metro Atlanta

Species richness between dry-air and rain samples, based on the Chao1 nonparametric metric (Chao, 1984) (Kim et al., 2017), appeared to be similar for both bacteria and fungi (Figure 1A, 1E) as determined by ANOVA analysis (p-value > 0.18). Species richness represents the number (or count) of individual species in a community, without taking into account relative abundances. This result was also observed for the Pielou’s index, which quantifies how equal (even) a community of different species numerically is. The calculation of Pielou’s index suggests that the atmosphere overall has a microbial richness that appeared to be relatively consistent between rain and dry-air samples over the two years of sampling.

**Figure 1.**
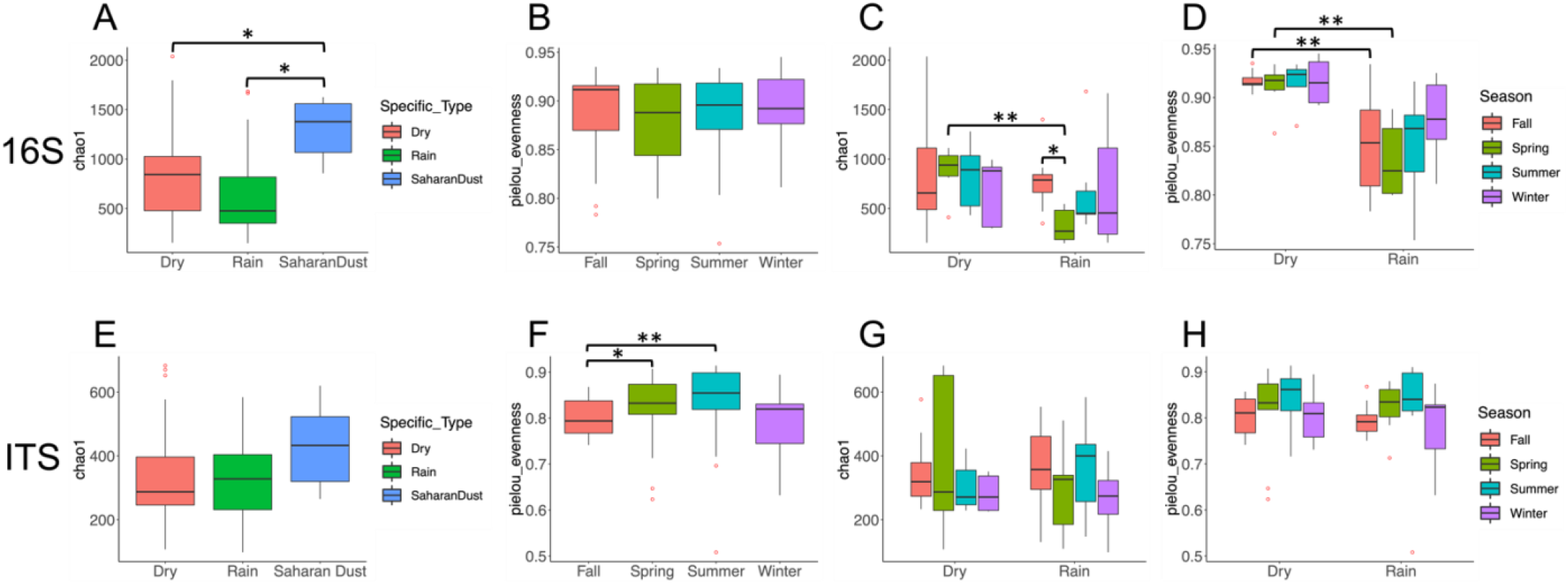
Microbial diversity patterns for bacteria (16S, top) and fungi (ITS, bottom) by sample type and seasonality. (A) Chao1 bacterial species richness, which details the number of individual species in a community, without taking into account relative abundances, for each sample type, i.e., dry, rain and Saharan dust. (B) Pielou bacterial species evenness, which quantifies how equal (even) a community of different species is, by the seasons. (C) Bacterial richness by seasonality and sample type (i.e., dry and rain). (D) Bacterial evenness by seasonality and sample type (i.e., dry and rain). (E) Fungal richness by sample type (i.e., dry, rain and Saharan dust). (F) Fungal evenness by seasons. (G) Fungal richness by seasonality and sample type (i.e., dry and rain). (H) Fungal evenness by seasonality and sample type (dry and rain).

### Taxonomic differences between dry-air and rain samples

The taxonomic profiles of the field samples revealed that the *Methylobacterium-Methylorubrum*, *Massilia*, *Pseudomonas*, *Hymenobacter*, *Deinococcus,* and *Pantoea* were the most abundant genera in all sample types overall (Figure 2) and showed similar relative abundances between dry-air and rain samples. For the fungal community, the most abundant genera were *Alternaria*, *Cladosporium*, *Trametes*, and *Epicoccum* (Figure 2) which showed clear preferences for specific sample types, discussed below.

**Figure 2.**
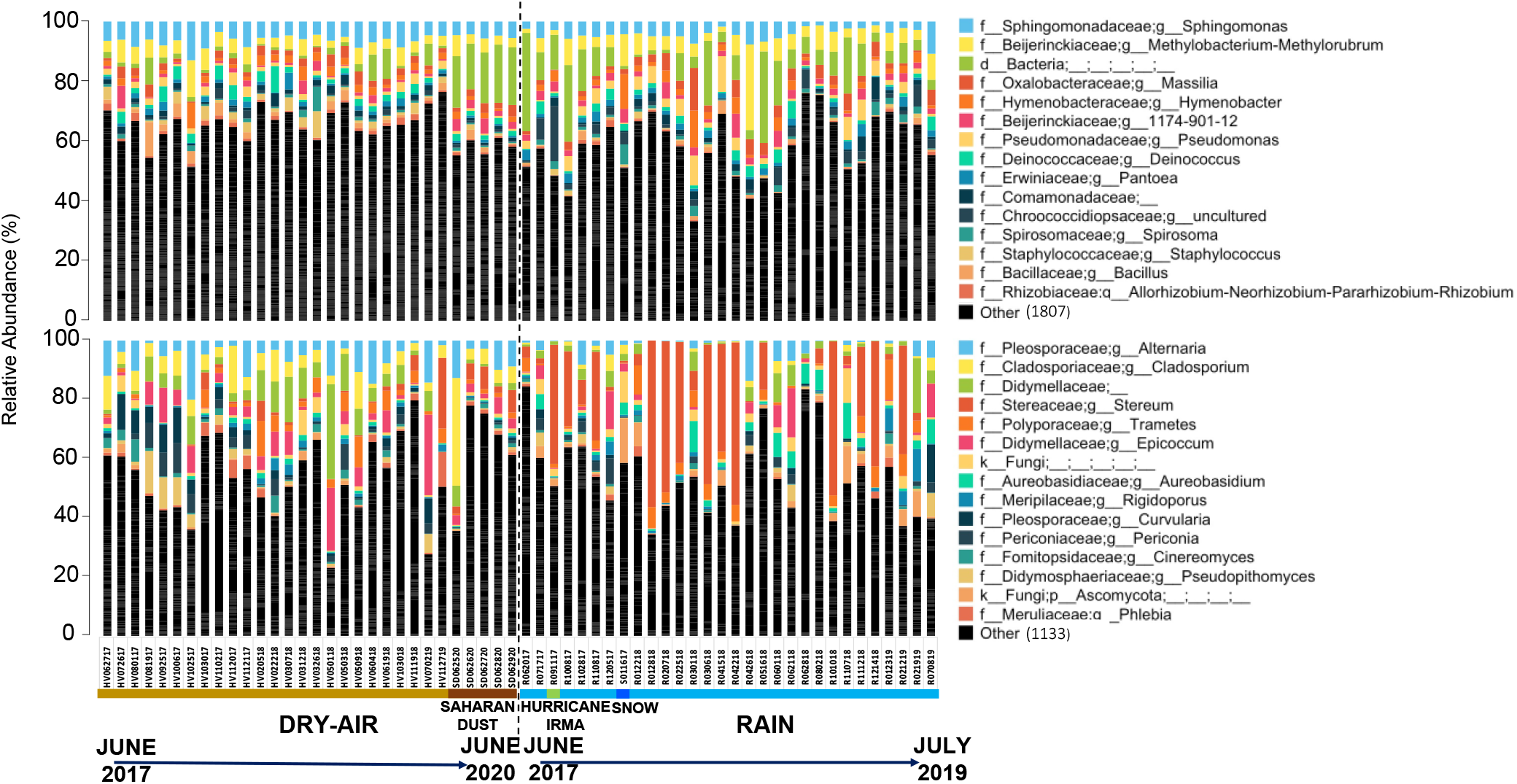
Bacterial and fungal taxonomic profile for dry-air, rain, and Saharan dust samples at genus level. The relative abundance was determined as described in the Materials and Methods section and is represented by the size of the bar plot (see also Figure key for taxon identification).

Examining differential abundance between sample types based on the Lefse tool and the phylum level, *Proteobacteria* [recently proposed to be renamed as *Pseudomonadota* (Oren & Garrity, 2021) was the most abundant phylum across the whole sample set but was significantly more abundant in rain samples by Lefse (p-value ≤ 0.05). *Proteobacteria* relative abundance ranged from 27% to 72%, depending on the sample considered, followed by the *Cyanobacteria* and *Abditibacteroita* phyla (Fig. S9). On the other hand, dry-air samples showed an enrichment in *Firmicutes (Bacillota), Actinobacteriota (Actinomycetota)*, *Chloroflexi (Chloroflexota),* and *Planctomycetes (Planctomycetota)* bacterial phyla and *Halobacterota, Euryarchaeota*, and *Crenarchaeota* archaeal phyla (p-value ≤ 0.05) (Fig. S9). In the fungal dataset, nine different phyla were identified in our ITS dataset, and none were found to be significantly higher in abundance in a specific sample type (p-value ≥0.05). The most abundant phyla across all samples were *Basidiomycota* and *Ascomycota*.

Furthermore, gram-positive and spore-forming bacterial families such as *Staphylococcaceae* (Fig. S10B), including the *Staphylococcus* and *Salinicoccus* genera (Fig. S11B), and the *Bacillaceae* family, including the *Bacillus* genus, appeared to be higher in dry-air samples (p-value ≤ 0.05), which is consistent with previous bioaerosols studies (Be et al., 2015). Gram-negative bacteria with diverse metabolic capabilities such as members of the *Sphingomonadaceae* family (Fig S10B), including the *Sphingomonas* genus (Fig S11B), and those related to plant growth enhancement, such as *Rhizobiaceae* (Fig. S10B), were also more abundant in dry-air by Lefse (p-value ≤ 0.05). The most abundant fungi in dry-air samples were related to plant pathogens and widespread fungal families such as *Pleosporaceae, Didymellaceae* including *Epicoccum, Cladosporiacea* (Fig. S12A), and two common indoor and outdoor airborne fungi*— Cladosporium* and *Toxicocladosporium* (p-value ≤ 0.05) (Fig. S13B). Other significantly enriched fungal genera in dry-air samples were related to plant pathogen members of the *Nigrospora* genus (p-value ≤ 0.05) (Fig. S13B).

On the other hand, photosynthetic bacterial families such as *Chroococcidiopsaceae, Leptolyngbyaceae* (including the *Leptolyngbya* genus), and unclassified members of the *Cyanobacteria* class were higher in abundance in rain samples (Fig. S10B). Members of the *Erwiniaceae* family, which includes plant pathogens, insect endosymbionts, and some representatives with known ice-nucleating capabilities, were also more abundant in rain samples. In the case of fungi, wood decaying fungal families such as *Stereaceae* (including *Stereum*) and *Diatrypaceae,* and the cosmopolitan family *Aureobasidiaceae* were higher in abundance in rain samples (Fig. S12B). In addition, saprotrophic fungi (*Exidia*) and plant pathogens on grain crops *(Ustilago*) were also higher in abundance in rain samples (Fig. S13B). Finally, to further assess the effects of sample type (dry-air vs. rain), non-metric multidimensional scaling (NMDS) visualization showed that bacterial communities were more well-defined by sample type than fungal communities (Figure 3A-3B).

**Figure 3.**
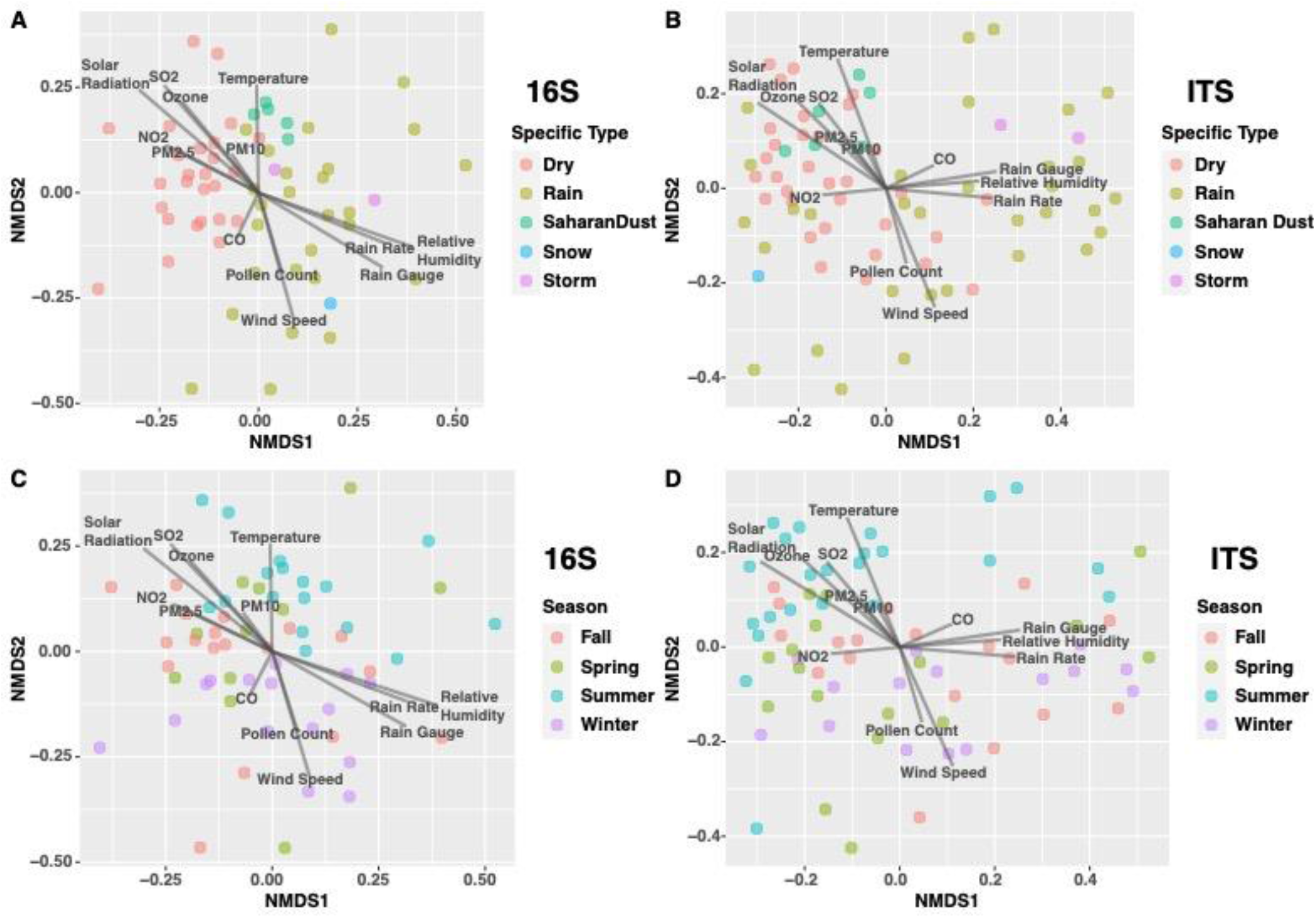
Effect of environmental factors on airborne bacterial and fungal community similarity. The graph represents the non-metric multidimensional scaling (NMDS) plot based on the Bray-Curtis distance matrix of bacterial and fungal communities in all samples used in the study. Note the significant effect of sample type. For bacteria, stress value = 0.2150442, for fungi stress value = 0.1815254.

### Dry samples have higher relative abundance of pathogenic fungi

We identified several known pathogens of humans and plants in the fungal dataset based on classification of ASVs at the species level. *Alternaria alternata,* which is known to cause leaf spot on several plants and upper respiratory infections to immunocompromised humans (Gabriel et al., 2016) was identified in most samples, except for a few rain samples. *A. alternata* was significantly higher in abundance in dry-air (5.6% compared to 2.3% in rain samples; p-value = 0.001 with ANOVA analysis). Another pathogenic fungal species identified in our samples was *Cladosporium cladosporioides* with a relative abundance of 2.2% in dry-air vs. 0.7% in rain (p-value = 0.0001). *C. cladosporioides* is a saprobic fungi that is common in indoor and outdoor settings and an opportunistic allergen (Bensch et al., 2010). Moreover, a few bacterial genera that include pathogenic members, including *Bacillus* and *Staphylococcus*, were identified in dry-air, rain, and Saharan dust samples. Average relative abundance of these bacterial genera all together was 8.5% and 6.5% for dry-air and rain, respectively (Fig. S14). In addition, our data had enough resolution to identify pathogenic species at the ASV level in some cases, including *Legionella pneumophila*, *Bacillus halodurans*, *Clostridium perfrigens*, *Campylobacter jejuni*, *and Corynebacterium glutamicum,* but the abundance of these species did not significantly differ between dry-air and rain samples and was always low e.g., <0.01% of total.

### Seasonality did not have a strong effect on microbial diversity and composition in Metro Atlanta

Our results showed that seasonality did not have a strong effect on microbial richness, independent of the type of sample evaluated (Fig. S15). It should be mentioned, however, that fungi had less species evenness in fall compared to spring (p-value =0.04, by Krustal-Wallis, as implemented in QIIME2), and summer, (p-value = 0.008) (Figure 1F), when dry-air and rain sample types were evaluated together. Further, the seasonality effect on the bacterial dataset appeared to be significant when the analysis was performed separately by sample type, but not when combined. Specifically, bacterial species richness appeared to be higher in the fall compared to the spring season (p-value = 0.02) only in rain samples (Figure 1C). Notably, the driest months (lowest average precipitation) in our sampling region are between September and November (fall season), based on data collected over 26 years (1996 – 2021). In addition, our analysis showed higher bacterial species richness in dry-air than in rain samples only during the spring season (p-value = 0.003) (Figure 1C), and higher bacterial evenness in dry-air than in rain samples during both the spring (p-value = 0.005) and the fall (p-value = 0.002) seasons (Figure 1D). Collectively, these results revealed minor differences in richness between the seasons with significant differences observed when assessing richness within one sample type over the year, or by evaluating diversity between dry-air and rain samples in a specific season.

Moreover, by evaluating NMDS plots of the Bray-Curtis distance matrix of the samples, a notable overlap was observed mostly between fall, spring, and summer seasons for bacteria while winter appeared to be less similar and more scattered (Figure 3C). Fungal community composition appeared to be more randomly mixed between all seasons, except for the summer season, when the fungal community compositions appeared to be more tightly clustered (Figure 3D). Relative abundance comparison between seasons at class level for the bacterial and fungal datasets did not reveal any significant trend or difference between seasons (Fig. S16).

### Variance in bacterial and fungal communities is mostly explained by sample type

To evaluate the variables that drove the microbial community compositional differences observed between samples, we used the Adonis function within the vegan package in RStudio. The analysis showed that bacterial and fungal communities were mostly influenced by sample type (R^2^ = 12.3% and 15.5%, respectively), followed by seasonality (R^2^ = 6.8% and 10.3%, respectively) with a p-value = 0.001 (Table 1 and 2). Other variables analyzed such as back trajectory (air mass origin), land use (Table 1 and 2), air pollutants (O_3_, NO_2_, CO, and SO_2_), and particulate matter (PM2.5 and PM10) (Table S2 and S3) did not appear to significantly explain differences. In addition, the variability in measured DNA concentration and number of sequences obtained did not appear to explain the variance between samples (Table 1 and 2). Thus, around 78% of the variance is presumably attributed to other, not measured variables/conditions (Table 1 and 2) and/or general variabilities in the composition of the lower troposphere in Atlanta.

**Table 1.**
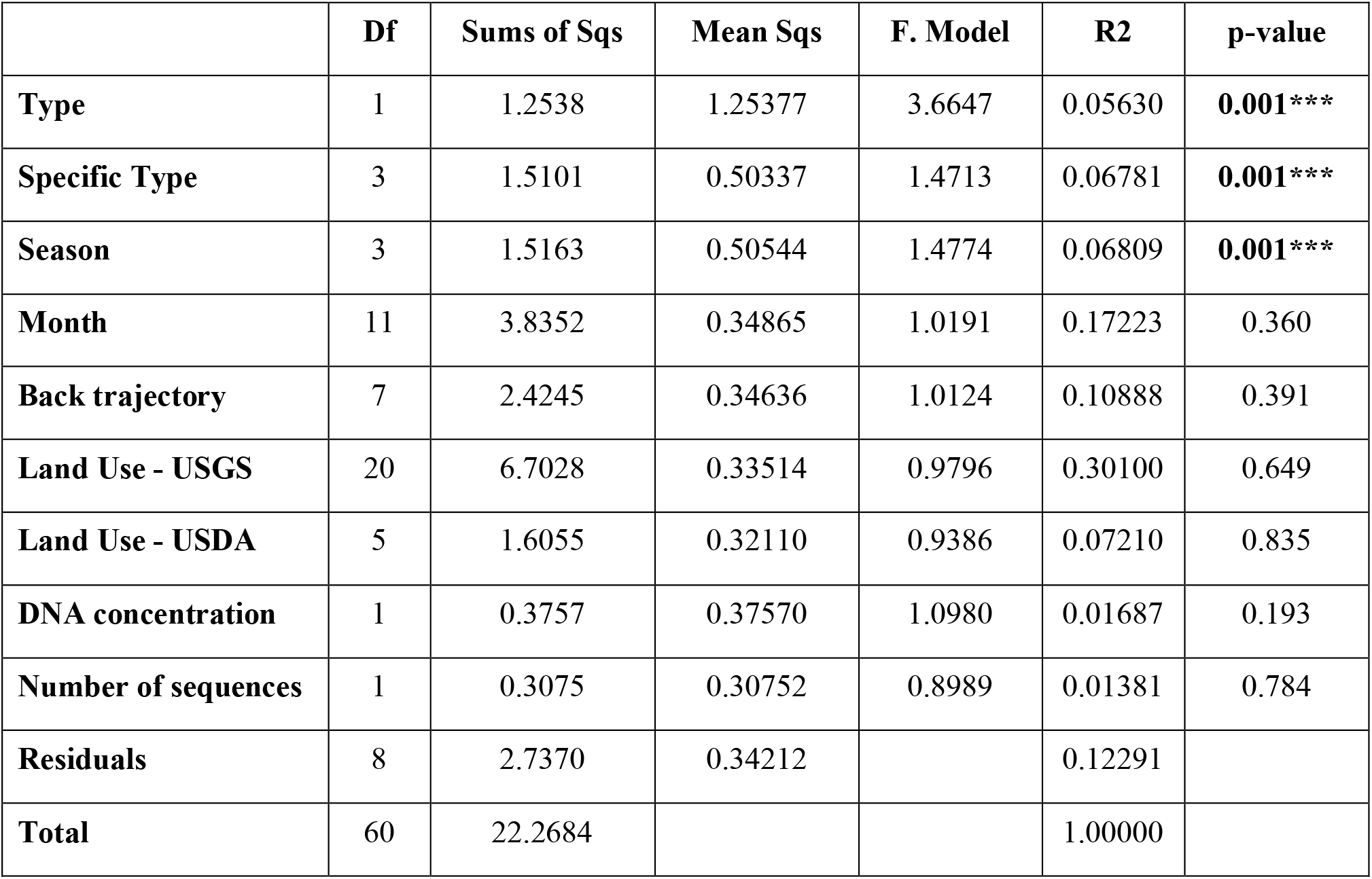
Adonis analysis of 16S rRNA gene amplicon data. Significance is denoted by asterisks.

|  | **Df** | **Sums of Sqs** | **Mean Sqs** | **F. Model** | **R2** | **p-value** |
| --- | --- | --- | --- | --- | --- | --- |
| **Type** | 1 | 1.2538 | 1.25377 | 3.6647 | 0.05630 | **0.001\*\*\*** |
| **Specific Type** | 3 | 1.5101 | 0.50337 | 1.4713 | 0.06781 | **0.001\*\*\*** |
| **Season** | 3 | 1.5163 | 0.50544 | 1.4774 | 0.06809 | **0.001\*\*\*** |
| **Month** | 11 | 3.8352 | 0.34865 | 1.0191 | 0.17223 | 0.360 |
| **Back trajectory** | 7 | 2.4245 | 0.34636 | 1.0124 | 0.10888 | 0.391 |
| **Land Use - USGS** | 20 | 6.7028 | 0.33514 | 0.9796 | 0.30100 | 0.649 |
| **Land Use - USDA** | 5 | 1.6055 | 0.32110 | 0.9386 | 0.07210 | 0.835 |
| **DNA concentration** | 1 | 0.3757 | 0.37570 | 1.0980 | 0.01687 | 0.193 |
| **Number of sequences** | 1 | 0.3075 | 0.30752 | 0.8989 | 0.01381 | 0.784 |
| **Residuals** | 8 | 2.7370 | 0.34212 |  | 0.12291 |  |
| **Total** | 60 | 22.2684 |  |  | 1.00000 |  |

**Table 2.** Adonis analysis of ITS amplicon data. Significance is denoted by asterisks.

|  | **Df** | **Sums of Sqs** | **Mean Sqs** | **F. Model** | **R2** | **p-value** |
| --- | --- | --- | --- | --- | --- | --- |
| **Type** | 1 | 2.2565 | 2.25654 | 8.2286 | 0.09352 | **0.001\*\*\*** |
| **Specific Type** | 3 | 1.5005 | 0.50016 | 1.8239 | 0.06219 | **0.001\*\*\*** |
| **Season** | 3 | 2.4894 | 0.82980 | 3.0259 | 0.10317 | **0.001\*\*\*** |
| **Month** | 11 | 3.5533 | 0.32303 | 1.1779 | 0.14727 | 0.077 |
| **Back trajectory** | 7 | 1.9672 | 0.28103 | 1.0248 | 0.08153 | 0.398 |
| **Land Use - USGS** | 20 | 5.0964 | 0.25482 | 0.9292 | 0.21122 | 0.783 |
| **Land Use - USDA** | 5 | 1.5318 | 0.30635 | 1.1171 | 0.06348 | 0.204 |
| **DNA concentration** | 1 | 0.2537 | 0.25370 | 0.9251 | 0.01051 | 0.538 |
| **Number of sequences** | 1 | 0.2694 | 0.26939 | 0.9823 | 0.01116 | 0.441 |
| **Residuals** | 19 | 5.2104 | 0.27423 |  | 0.21594 |  |
| **Total** | 71 | 24.1286 |  |  | 1.00000 |  |

### Saharan dust carries pathogenic bacteria and fungi to Metro Atlanta

In this study, we classified three major weather events as foreign atmospheric events: the rain sample from the remnants of Hurricane Irma in 2017, dry-air samples from a Saharan dust cloud event in 2020, and a rare snow event in 2018. Rain samples collected during the Hurricane Irma remnants and the 2018 snow event sampling showed an enrichment of *Cyanobacteria* (relative abundance of 26.2% and 17.5%, respectively) compared to the average relative abundance (5.7%) in the typical rain events in Atlanta. Saharan dust showed an enrichment of unclassified *Bacteria*, *Myxococcota*, and uncultured SAR324_clade (Marine_group_B), the latter being consistent with oceanic transport of air masses (Fig. S9). The Hurricane Irma rain sample showed an enrichment of *Agaricomycetes* (68.6%) and decreased abundance of *Dothideomycetes* (12.1%) compared to the remaining rain samples (53% and 26%, respectively). Saharan dust samples appeared to have transported relatively more cells belonging to *Agaricomycetes* as evidenced by the taxon’s relative abundance between sample SD062520, taken the day before the dust cloud arrived (16% relative abundance) and the samples taken for the following four days that the dust cloud was moving out of our sampling area (samples SD062620, SD062720, SD062820, SD062920, SD070220 with 81.2%, 78.5%, 66.4%, 63.2%, and 46.5% relative abundance, respectively). *Agaricomycetes* are a mushroom-forming soil fungal class, and some of its species are pathogenic. Furthermore, the air masses sampled during the Sahara dust event had a higher abundance of potential human pathogens affiliated with the bacterial *Rickettsiaceae* family (S10A) as well as the fungal *Alternaria alternata* (8.3%) and *Cladosporium cladosporioides* (3.7%) when compared to dry-air and rain samples with p-values = 0.002 and 0.00007, respectively by ANOVA. However, other important fungal pathogens such as *Cryptococcus neoformans* and *C. gatti*, the causative agents of cryptococcal meningitis often associated with sub-Saharan Africa aerosols (Assogba et al., 2015), as well as *Histoplasma* and *Pneumocystis* fungi, were not identified in our dataset. Foreign air masses brought by the Saharan dust event also appeared to carry a higher bacterial species richness compared to the average dry-air (p-value = 0.03) and rain samples (p-value = 0.01) (Figure 1A). Fungal species richness did not appear to be significantly affected by the Sahara event overall (Figure 1E).

## Discussion

The dry-air and rainfall samples collected over two years showed that Metro Atlanta, as an urban and heavily populated setting, has a relatively consistent microbial richness (bacteria and fungi), with little seasonality effects. Moreover, our results indicated that there is no difference in the bacterial and fungal diversity (species richness and evenness) between dry-air samples and rain samples over a period of two years. These results contrast with those of a previous study performed at high elevation (2896 m asl) in Sierra Nevada, Spain that found that bacterial species richness was significantly higher in dry deposition (Triadó-Margarit et al., 2019). Given also that our samples represent robust measurements over a two-year period, these results indicate that the patterns of airborne microbial diversity may differ substantially between regions.

Dry days (dry-air samples) appear to promote the enrichment of spore-forming bacteria (*Bacillus*), gram-positive bacteria (*Firmicutes*; *Staphylococcus*), plant pathogens, widespread fungi (*Pleosporaceae, Didymellaceae*), and common airborne fungi (*Cladosporium*) that generally were not present in rain samples. In contrast, photosynthetic bacteria (*Cyanobacteria)*, plant pathogens and potential IN bacteria (*Erwiniaceae*), unclassified *Bacteria*, and wood decaying fungi (*Stereum*) were enriched in rain samples. Previous literature suggests that rainfall is not necessarily a washout of all the microbes that are suspended in the atmosphere (Els et al., 2019). Here we report that rain appears to carry a significant abundance of habitat-specific microbes (hydrophilic, aquatic bacteria and fungi related to wood-decay). In agreement with our results, significant differences in terms of taxonomic composition have been reported for fungi in wet deposition, including the enrichment of *Stereum* species, compared to dry deposition, suggesting the presence of fungal taxa that are deposited primarily with rain (Woo et al., 2018). In addition, the enrichment of *Cyanobacteria* in the atmosphere during rain events has been reported previously (Seifried et al., 2015; Dillon et al., 2020; Wiśniewska et al., 2022). Similar results showing differences between the microbial composition in dry air and rain have only been reported at high-elevation study sites (Els et al., 2019; Triadó-Margarit et al., 2019).

Several of the taxa reported to be enriched in rain samples match those reported here, but others do not. Most notably, *Cyanobacteria* and *Stereaceae* were not enriched in rain samples at high elevations, unlike in our study. These results suggests that local microbial sources might influence or shape the microbial community in rain. Instead, *Actinobacteria, Acidobacteria, Noviherbaspirillum, and Massilia* were enriched in rain samples at high elevations and remote areas (Els et al., 2019; Triadó-Margarit et al., 2019) but not in our samples. In addition, previous results on the decreased abundance of *Cyanobacteria* in aerosol samples after rainfall were based on samples collected adjacent to marine environments (Baltic Sea), and only for a couple of months during the heavy rain season and with culture-dependent techniques (Wiśniewska et al., 2022). Therefore, our results highlight the importance of studying the airborne microbial communities in different regions and conditions, e.g. urban and low-altitude studied here, and for longer periods of time (year-round). Cities are important modern sources of bioaerosols caused by anthropogenic activities. For instance, in Japan, bacteria associated with human skin such as *Propionibacterium*, *Staphylococcus*, and *Corynebacterium* were higher in abundance in an urban site while an enrichment of *Methylobacterium* and *Sphingomonas*, which are mostly associated with soil and plants, was observed in a suburban site (Tanaka et al., 2020).

Our study showed the clear enrichment of pathogenic fungi in dry-air relative to rain samples. A previous study reported similar findings for pathogenic fungi in urban spaces (Niu et al., 2021). However, the opposite pattern had been reported in forested spaces, where pathogenic microbes and allergens were more abundant in air samples during rain events (Huffman et al., 2013). It should be mentioned, however, that the two studies mentioned above only sample air bioaerosols when collecting rain data, whereas our study directly collected rain samples. Thus, our study contributes new insights into the potential differences of microbial diversity and composition in dry-air and rain samples, and highlight that aerosolized microbes following rain events may differ significantly from microbes deposited with the rain. The presence of potential pathogenic bacteria and fungi has previously been reported in air samples collected in urban settings (Nicolaisen et al., 2017; Franchitti et al., 2022), but our findings shed light on previously unknown comparisons between dry and rain conditions in these environments, with implications for public health since our results show that dry air samples represent likely a higher health risk due to the enrichment of pathogens and allergens.

In terms of seasonal patterns, our richness results contradict those published previously for airborne microbes (Bowers et al., 2013; Els et al., 2019). Bacterial and fungal species richness in the collected samples were relatively constant across seasons during the two years of sampling. In the study by Els and colleagues (2019), August (summer) samples had the lowest bacterial species richness, while the summer season in our study showed the highest species richness for both bacteria and fungi. In the same study, bacterial species richness was highest in November while fungal species richness was lowest in May. Our lowest species richness for both bacteria and fungi appeared during the winter season. The notable differences between our study and the Els et al. (2019) study may be attributed to collection site differences. The Els study was conducted above the planetary boundary layer in Mount Sonnblick (3106 m asl in the Austrian Alps), which is a remote site with little anthropogenic activity. In addition, seasonality was represented by only one month per season while our study collected representative samples from all months for the four seasons. Thus, the differences observed highlight the importance of consistent, long-term, and local bioaerosols studies, especially in urban settings where the diversity of the aerosolized microbes is likely higher (Ruiz-Gil et al., 2020) and potentially more dynamic, with wider sources of potential or opportunistic pathogens.

Although the effect of seasonality on bacterial diversity patterns showed some weak significance when rain samples were analyzed independently of dry samples, or when all samples from one season were compared, the effect was not significant when dry days and rain were analyzed together across the whole year. The lack of a clear seasonality pattern in our study contradicts previous studies that had identified seasonal patterns in the airborne microbial composition (Bowers et al., 2012, 2013; Be et al., 2015; Uetake et al., 2019; Els et al., 2019; Núñez et al., 2021). The weakly significant seasonality effect in our datasets could reflect the conditions of a humid subtropical climate. The climate of Atlanta has short, mild winters and long, hot, and humid summers. In contrast, previous studies conducted at a high-elevation site in Colorado, USA, observed a stronger seasonality effect, driven by more dramatic temperature changes and snow events (Bowers et al., 2012). Temperature varies less between seasons in Atlanta, and snow events are rare. Consistent with these interpretations, the bacterial taxa related to cold environments such as the *Moraxellaceae* family were significantly higher in abundance in the cold and snowy months (November - April) compared to the warmer months in Colorado, while our study did not note the same significant difference in taxa related to cold environments between seasons. The strong seasonal pattern in chloroplast sequences during the early spring (blooming season) captured by our study indicated that our methodology was robust. Therefore, despite disagreements with past studies, we are confident in our conclusion that seasonality plays a minimal role in Metro Atlanta microbial diversity. In addition, several previous studies that reported strong seasonal patterns pooled samples from different days or weeks (Be et al., 2015; Mu et al., 2020), whereas no sample pooling was performed in our study. Further studies, as well as a consensus on sampling instruments, techniques, and sample processing, will be needed to further quantify the seasonal patterns in airborne microbes in the Southeast US and elsewhere. Additionally, the importance of local environmental conditions and bioaerosol sources should not be underemphasized.

Interestingly, the Saharan dust samples had the highest bacterial richness compared to local dry-air and rain collected at Metro Atlanta. Similar results for Saharan dust intrusion events have been reported previously in the Iberian Peninsula (González-Toril et al., 2020). Furthermore, the presence of Saharan dust in the Metro Atlanta area, as an uncommon and foreign bacterial and fungal source, appeared to substantially alter the usual bacterial and fungal communities, with the notable enrichment of potential human bacterial pathogens (*Rickettsiaceae*) and pathogenic fungi and fungal allergens (*Alternaria alternata* and *Cladosporium cladosporioides*). Therefore, our results collectively indicate that major Saharan dust clouds could potentially have public health consequences, especially for people with a compromised respiratory system, consistent with the results of previous studies (de Longueville et al., 2013; Zhang et al., 2016; Marone et al., 2020).

Our results demonstrate that bacterial and fungal communities in rain differ from those in dry air, suggesting that different sample types should be targeted for a better understanding the atmosphere as a microbial habitat. Our study also suggests that a strong seasonal pattern for airborne microbes does not exist in an urban setting in Southeast USA, and indicates that dry-air samples and samples from Sahara dust may contain a higher relative abundance of pathogens and allergens. More intensive sampling during the seasons coupled to higher-resolution, genome-resolved methods (metagenomics) will be needed to better understand seasonality patterns for urban settings or cities and more precisely identify pathogens compared to the amplicon approach employed here that typically cannot provide strain-level resolution. The high complexity and heterogenicity of the atmosphere and variety of local sources makes it challenging to establish clear microbial patterns and the effects of environmental factors on these patterns. Despite these challenges, however, our study provides important novel insights into the significance of dry vs. rain sampling methods on assessing the composition and diversity of microbial communities in a given environment. Additionally, our study emphasizes the impact of weather patterns, pollutants, and air masses on microbial community composition, while providing findings that are generally consistent with what has been previously reported.

## Acknowledgements

This work was supported in part by a NASA Exobiology program (award no. 20-EXO20-0007) and from NSF Dimensions of Biodiversity program (award no. 1831582) to KTK.

